# Direct Nucleosome Binding of Borealin Secures Chromosome Association and Function of the Chromosomal Passenger complex

**DOI:** 10.1101/620385

**Authors:** M. A. Abad, J. G. Ruppert, L. Buzuk, M. Wear, J. Zou, K. M. Webb, D. A. Kelly, P. Voigt, J. Rappsilber, W. C. Earnshaw, A. A. Jeyaprakash

**Affiliations:** Wellcome Centre for Cell Biology, University of Edinburgh, Edinburgh EH93BF, UK; Technical University, Berlin 13355, Germany

## Abstract

Chromosome association of the Chromosomal Passenger Complex (CPC; consisting of Borealin, Survivin, INCENP and the Aurora B kinase) is essential to achieve error-free chromosome segregation during cell division. Hence, understanding the mechanisms driving the chromosome association of the CPC is of paramount importance. Here using a multifaceted approach, we show that the CPC binds nucleosomes through a multivalent interaction predominantly involving Borealin. Strikingly, Survivin, previously suggested to target the CPC to centromeres [1–3] failed to bind nucleosomes on its own and requires Borealin and INCENP for its binding. Disrupting Borealin-nucleosome interactions excluded the CPC from chromosomes and caused chromosome congression defects. We also show that Borealin-mediated chromosome association of the CPC is critical for Haspin- and Bub1-mediated centromere enrichment of the CPC and works upstream of the latter. Our work thus establishes Borealin as a master regulator determining the chromosome association and function of the CPC.

## Introduction

Chromosome segregation is a complex process involving numerous protein-protein and protein-DNA interactions tightly controlled by signaling networks consisting of kinases and phosphatases [4–6]. Aurora B kinase, the enzymatic core of the Chromosomal Passenger Complex (CPC) is a key regulator essential for error-free chromosome segregation and functions by controlling multiple steps of cell division: chromosome condensation and cohesion, kinetochore-microtubule attachments, the spindle assembly checkpoint and cytokinesis (reviewed in [4, 7, 8]). The CPC is composed of Aurora B, INCENP, Borealin/Dasra and Survivin and can be divided into distinct localisation and kinase modules, linked by the central helical coil of INCENP. The localisation module (CPC_LM) consisting of Borealin, Survivin and the first 58 amino acids (aa) of INCENP controls the localisation of the CPC [9, 10]. The kinase module consists of Aurora B and the IN-box of INCENP, a well-conserved C-terminal region required for full activation of Aurora B kinase [11–13]. CPC function is tightly linked to its distinct localisation during different stages of cell division. During early stages of mitosis the CPC localises to chromosome arms where it influences chromosome condensation [14, 15]. It subsequently concentrates at the inner centromere where it releases incorrect attachments and regulates the timing of mitotic progression via the spindle assembly checkpoint [8]. During anaphase, the CPC associates with the central spindle and during cytokinesis with the equatorial cortex and midbody to control cell abscission [16–20].

CPC localisation at centromeres has been suggested to depend on the co-existence of two histone modifications: Haspin-mediated phosphorylation on histone H3 Thr3 (H3T3ph) and Bub1-mediated phosphorylation on histone H2A Thr120 (H2AT120ph). According to the proposed models, H3T3ph is directly recognised by the BIR domain of Survivin [1–3] and H2AT120ph is read by hSgo1, which then recruits Borealin [1, 21]. However, Haspin depletion by siRNA does not abolish CPC association to chromatin [3] and Survivin is not sufficient to achieve centromeric enrichment or chromosomal association of CPC in cells expressing Borealin lacking its C-terminal half [10]. These observations collectively highlight a central question that remains unanswered. How does the CPC associate with chromosomes during early prophase prior to its H3T3Ph-mediated centromeric enrichment?

## Results and Discussion

### Borealin nucleosome binding is essential for chromosome association of the CPC

Consistent with our previous observations (Jeyaprakash *et al.*, 2007), transient expression of a Myc-tagged Borealin lacking the first 10 aa and C-terminal half (Borealin 10-109) failed to rescue the siRNA-mediated depletion of endogenous Borealin and the CPC was completely excluded from chromosomes during the early stages of mitosis leading to chromosome congression defects (Figure 1A, B and S1A, B). This led us to hypothesise a direct role for Borealin in mediating CPC-chromosome interactions. To test this, we reconstituted nucleosome core particles (NCPs) containing homogenous H3T3ph modification *in vitro* and performed Electrophoretic Mobility Shift Assays (EMSAs) with recombinant CPC localisation module (CPC_LM: Borealin-Survivin-INCENP_1-58_, Figure S1C). The CPC_LM showed clear binding to modified NCPs as evidenced by its retarded mobility (Figure 1C). Interestingly, CPC_LM containing Borealin_10-109_ (CPC_LM_Bor10-109_) failed to interact with NCPs even when mixed at a 32 times molar excess (Figure 1C). This is particularly surprising, as several studies have shown previously that Survivin can bind synthetic N-terminal HH3 peptides phosphorylated at Thr3 through its BIR domain [1–3]. To test if Survivin on its own could bind H3T3ph tail in the context of NCPs, we analysed the binding of purified Survivin and H3T3ph NCPs by EMSA. Strikingly, Survivin did not bind H3T3ph NCPs (Figure 1D) possibly due to a lack of H3 tail accessibility within the NCP. Together, these data demonstrate Borealin to be a major contributor to CPC-nucleosome interactions. Considering Borealin’s direct role, we next asked if the CPC_LM can bind unmodified NCPs. Interestingly, the CPC retarded the mobility of unmodified NCPs in the EMSA assays confirming its binding to unmodified NCPs (Figure 1E). These observations establish that the CPC can bind NCPs in a H3T3ph-independent manner and the interaction is mainly mediated by Borealin.

**Figure 1.**
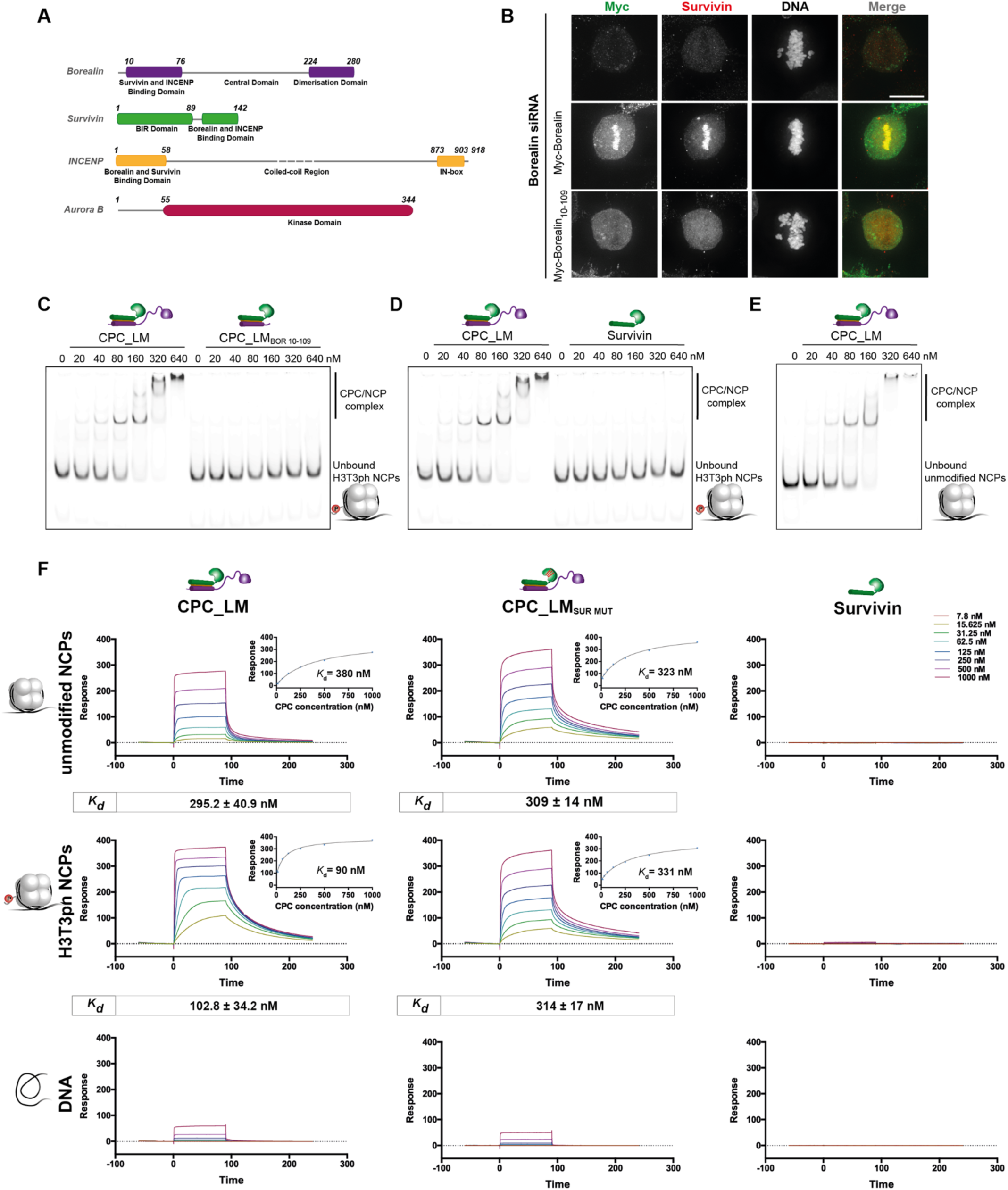
Borealin nucleosome binding is essential for chromosome association of the CPC. (A) Domain architecture of the subunits of the Chromosomal Passenger Complex (CPC). (B) Representative fluorescence images of a siRNA rescue assay for Borealin_10-109_ fragment. Immunofluorescent staining of Myc and Survivin in HeLa cells co-transfected with siRNA duplexes targeting the 3’UTR region of Borealin and Myc-Borealin constructs. Hoechst was used for DNA staining. Scale bar, 10 µm. All cells transfected with the siRNA and Myc-Borealin_10-109_ fragment showed exclusion of the CPC complex from the chromatin. (C,E) Native PAGE analysis of EMSA assays performed with increasing concentrations of recombinant CPC localisation module (CPC_LM) containing different Borealin fragments and either 20 nM phosphorylated (H3T3ph) (C) or unmodified IR700-labelled NCPs (E). (D) EMSA assays performed with increasing concentrations of Survivin with 20 nM IR700-labelled H3T3ph NCPs. (F) Representative SPR sensorgrams of the interaction between different CPC_LM complexes (CPC_LM, CPC_LM_SUR MUT_) or Survivin and unmodified (top) or H3T3ph NPCs (middle) or DNA (bottom) immobilized on the surface of a neutravidin sensor chip. Mean values (n ≥ 2, ± SEM) determined for the equilibrium dissociation constant (K*_d_*) are shown in boxes underneath the sensorgrams.

As H3T3ph has been proposed to be critical for concentrating the CPC at inner-centromeres, we speculated that phosphorylation on H3 Thr3 might positively influence nucleosome binding affinity of the CPC. To address this, we performed Surface Plasmon Resonance (SPR) experiments by flowing CPC at different concentrations over the sensor surface containing immobilised NCP and measured steady-state binding affinities. While the CPC_LM interacted with unmodified NCPs with a K_*d*_ of 295.2 ± 40.9 nM, interaction with H3T3ph NCPs was 3-fold tighter at 102.8 ± 34.2 nM (Figure 1F). Interestingly this increase in affinity for phosphorylated NCPs was due to Survivin binding to the H3 N-terminal tail, as a CPC containing a Survivin BIR mutant deficient for binding the phosphorylated H3 tail (CPC_LM_SUR MUT_, Figure S1D), bound both modified and unmodified NCPs with a similar affinity (Figure 1F). Thus, our data show that although the Survivin-H3 interaction is not essential for CPC chromatin binding per se, it enhances the affinity of the CPC for modified NCPs. Consistent with EMSAs, SPR failed to detect any binding of Survivin on its own to NCPs even when H3 is phosphorylated at Thr3 (Figure 1F).

### N-terminal 10 amino acids and C-terminal half of Borealin are required for CPC-chromatin interaction

Considering the essential contribution of Borealin towards nucleosome binding, we next mapped the regions of Borealin directly involved in nucleosome interaction. We reconstituted several versions of CPC_LM complexes containing different Borealin mutants (designed based on its domain architecture) (Figure 1A and S2A) and tested them in EMSA assays with and without H3T3 phosphorylation on NCPs (Figure 2A and S2B). Deleting a well-conserved unstructured central region of Borealin (aa residues 110-206, CPC_LM_BOR Δloop_, Figure S2C) abolished CPC binding to NCP almost completely. The deletion of either the N-terminal 10 (CPC_LM_BOR 10-end_) or C-terminal 59 aa of Borealin (CPC_LM_BOR 10-221_) also caused a noticeable reduction in binding (Figure 2A and S2B). Considering the qualitative nature of the EMSA assay, we evaluated the NCP-binding affinities of mutant CPC complexes in SPR assays (Figure 2B and S3A). Consistent with EMSA assays, CPC_LM_BOR 10-end_ showed 3-fold reduction in binding affinity as compared to CPC_LM. Both CPC_LM_BOR Δloop_ and CPC_LM_BOR 10-221_ exhibited even weaker NCP binding with measured affinities in the µm range (Figure 2B). Together our data show that the N-terminal 10 aa and C-terminal half of Borealin contribute to nucleosome binding, possibly through multiple physical contacts.

**Figure 2.**
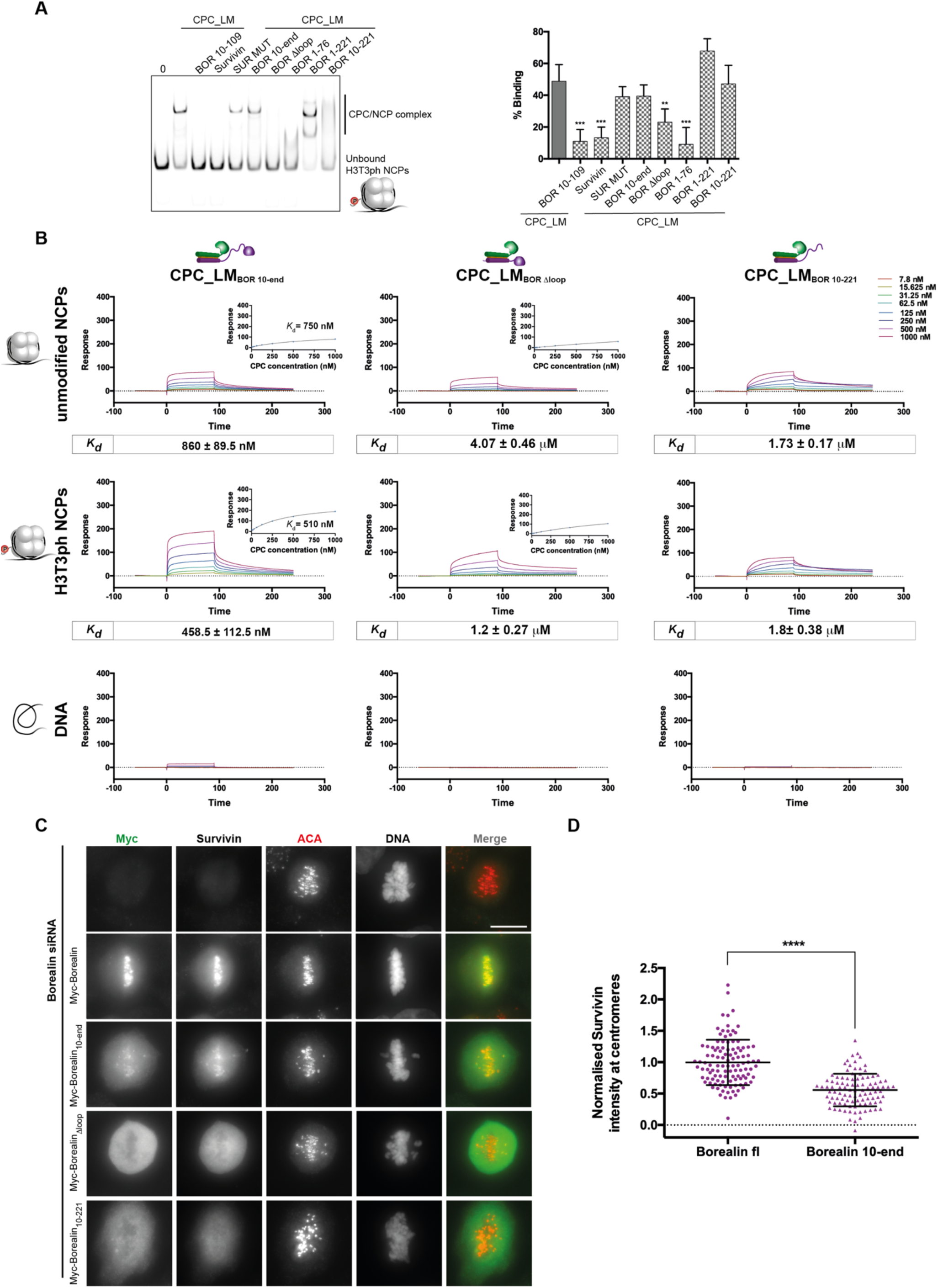
N-terminal 10 amino acids and C-terminal half of Borealin are required for CPC-chromatin interaction. (A) Native PAGE analysis of EMSA assays performed with recombinant CPC_LM containing different Borealin truncations binding to IR700-labelled H3T3ph NCPs (left) and quantification of binding (right). Concentrations of the NCP and the CPC used in the assay were 20 nM and 160 nM, respectively. Mean of % of binding ± standard deviation (SD); n=5; **P ≤ 0.01, *** P ≤ 0.001, unpaired *t*-test. (B) Representative SPR sensorgrams of the interaction between different CPC_LM complexes (CPC_LM_BOR 10-end_, CPC_LM_BOR Δloop_ and CPC_LM_BOR 10-221_) and unmodified (top) or H3T3ph NCPs (middle) or DNA (bottom). Mean values (n ≥ 3, ± SEM) determined for the equilibrium dissociation constant (K *_d_*) are shown in boxes underneath the sensorgrams. (C) Representative fluorescence images of a rescue assay for Borealin, Borealin_10-end_, Borealin_Δloop_, Borealin_10-221_ constructs. Immunofluorescent staining of Myc, Survivin and ACA in HeLa cells co-transfected with siRNA duplexes targeting the 3’UTR region of Borealin and Myc-Borealin constructs. Hoechst was used for DNA staining. Scale bar, 10 µm. (D) Quantification of Survivin levels at the centromeres for the siRNA-rescue assays with Myc-Borealin (n=111 cells) and Myc-Borealin_10-end_ (n=101 cells) shown in (C) (three independent experiments, mean ± SEM, Mann-Whitney test; P< 0.0001). For constructs Myc-Borealin_Δloop_ and Myc-Borealin_10-221_, all metaphase cells exhibited total exclusion of the CPC complex from the chromatin.

Having dissected the contribution of Borealin for nucleosome binding *in vitro*, we evaluated the behavior *in vivo* of Borealin mutants showing reduced NCP binding in siRNA rescue experiments (Figure 2C and S3B). Transient expression of Myc-Borealin_10-end_ in Borealin siRNA-depleted HeLa cells, resulted in a 50 % reduction of the centromeric association of the CPC (Figure 2D and S3C). In contrast, expression of Myc-Borealin_Δloop_ or Borealin_10-221_ resulted in almost complete exclusion of the CPC from chromosomes. Collectively, these observations demonstrate that Borealin-mediated nucleosome binding is essential for chromosome association of the CPC *in vivo*.

### CPC-Nucleosome binding is mediated by multivalent interactions predominantly involving Borealin

To gain structural insight into the underlying mechanism, the CPC-NCP complexes (Figure S4A) were crosslinked using EDC and analysed by mass spectrometry (Figure S4B, C). Consistent with our *in vitro* binding studies, regions of Borealin shown here to be critical for nucleosome binding made extensive contacts with NCPs, whereas Survivin interactions are mostly limited to the BIR domain and the H3 N-terminal tail (Figure 3A and S4D) as expected. Mapping the crosslinks onto the three dimensional structures of NCP and CPC (Figure 3B) suggests a model where the localisation module of the CPC together with a highly basic N-terminal tail of Borealin docks onto the acidic patch formed between H2A and H2B, a surface commonly involved in nucleosome recognition [22–26]. This interaction may orient the Survivin BIR domain to facilitate binding of Histone H3 tail phosphorylated at Thr3. Notably, Borealin loop residues trace along the DNA-histone interface implying its major contribution to nucleosome binding involves both protein-protein and possibly protein-DNA contacts.

**Figure 3.**
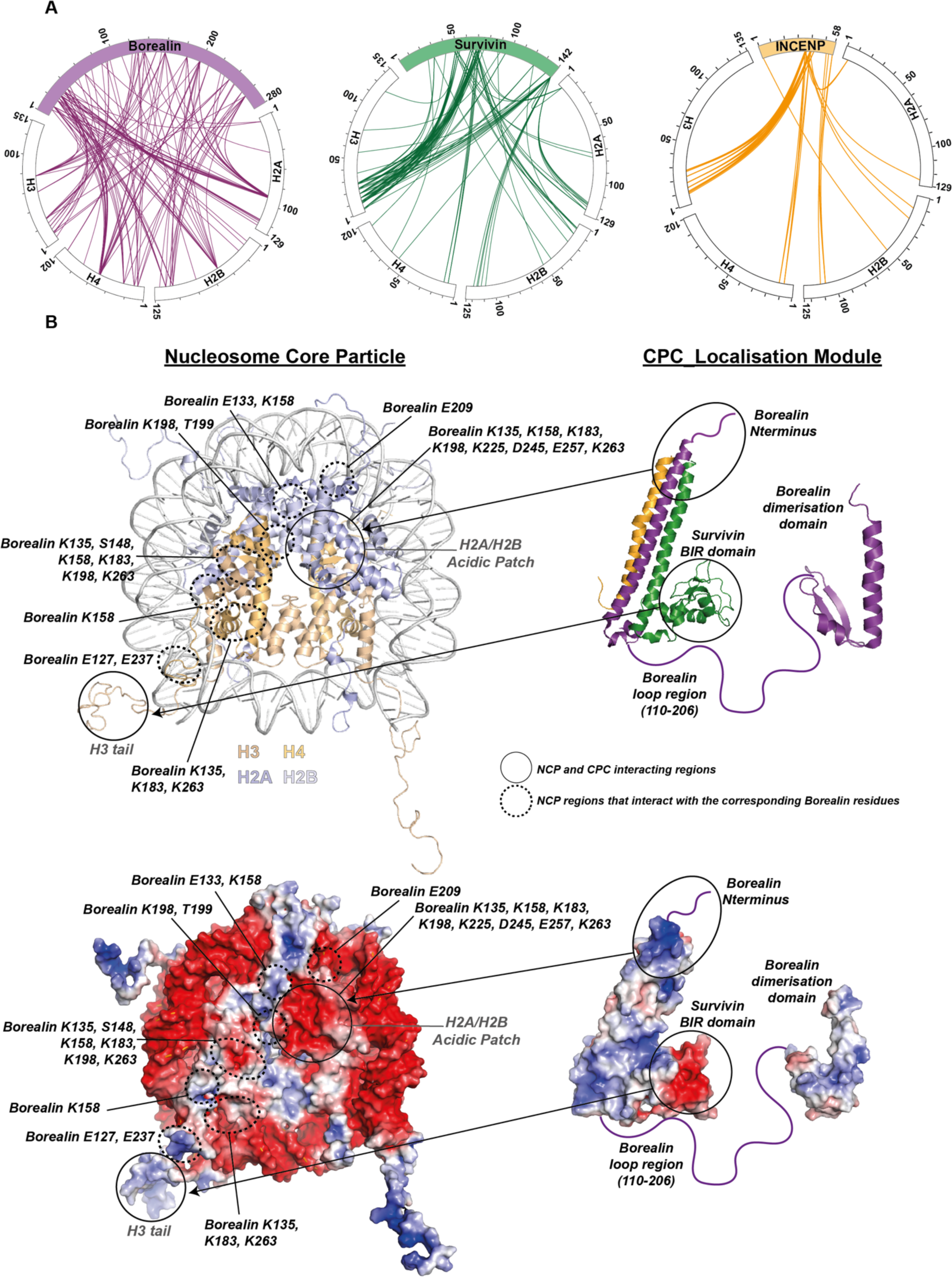
CPC-Nucleosome binding is mediated by multivalent interactions predominantly involving Borealin. (A) Circle view of the crosslinks between subunits of the CPC (Borealin-purple, Survivin-green, INCENP-yellow) and histones from unmodified NCPs. Intermolecular contacts involving Borealin, Survivin and INCENP and histones are shown as purple, green or yellow lines, respectively, using XiNET (Ref: PMID: 25648531). (B) Cartoon representation of the crystal/NMR structures of the NCP (PDB: 1kx5, [30]) and CPC (CPC core PDB: 2qfa, [10] and Borealin Dimerisation domain PDB: 2kdd, [31]) (top). Surface representation of the NCP and the CPC coloured based on the electrostatic surface potential calculated using APBS in Pymol v2.0.6 (bottom).

### Borealin-mediated chromosome association of the CPC is an upstream requirement for its Haspin and Bub1-mediated centromeric enrichment

As Haspin activity has been suggested to be stimulated by the Aurora B kinase, we next evaluated the impact of Borealin-mediated chromosome association of CPC on Haspin activity. Strikingly, in Borealin depleted cells, H3T3 phosphorylation was reduced to low levels (Figure 4A). While the expression of Myc-Borealin rescued these H3T3ph levels, expression of a Borealin mutant incapable of chromosome association (Myc-Borealin_Δloop_) failed to do so. Likewise, Borealin depletion led to a decrease in the levels of H2AT120 phosphorylation (Figure 4B), which could be rescued with Borealin, but only partially with Borealin_Δloop_. Notably, Haspin depletion, which resulted in reduced H3T3 phosphorylation and as a consequence diffused CPC localization along the chromosome arms, did not affect H2AT120 phosphorylation (Figure S5A). Together, our data demonstrates that CPC binding to nucleosomes is an upstream requirement for Haspin and Bub1 activities and for Haspin/Bub1-mediated CPC enrichment at centromeres.

**Figure 4.**
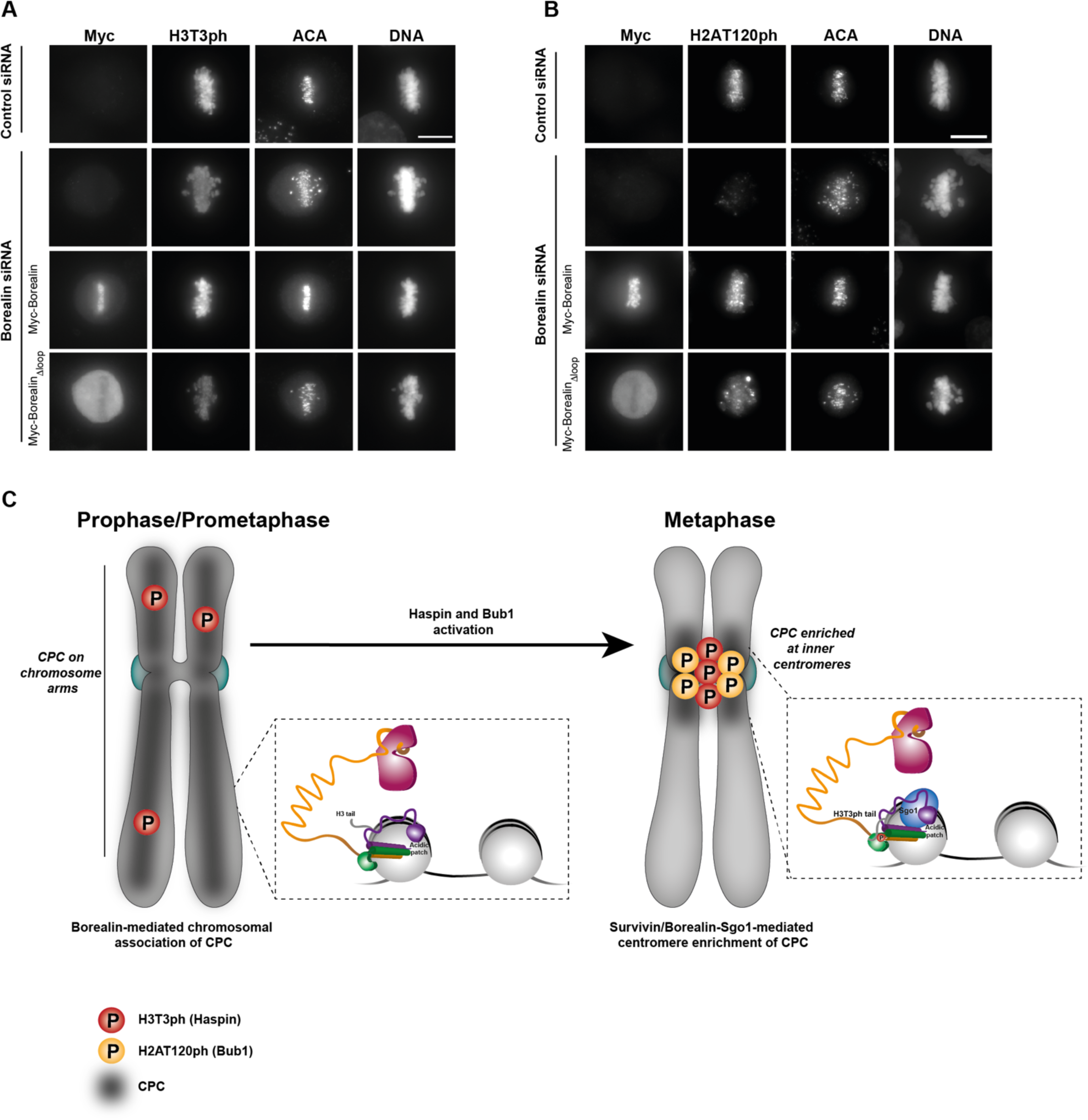
Borealin-mediated chromosome association of the CPC is an upstream requirement for its Haspin and Bub1-mediated centromeric enrichment. (A, B) Immunofluorescence analysis of ACA and H3T3ph (A) or H2AT120ph (B) levels upon Borealin depletion using siRNA duplexes and rescue with different Borealin constructs. Hoechst was used for DNA staining. Scale bar, 10 µm. (C) Model for Borealin-mediated chromatin association of the CPC and subsequent centromere enrichment.

In summary, our data suggest a mechanism for the chromosome association of CPC essential for error-free chromosome segregation (Figure 4C). During early mitosis when there is little or no H3T3ph [27] the CPC binds chromatin in a histone modification-independent manner, mainly via Borealin interactions involving multiple contacts with the histone octamer and possibly also with the DNA. Chromatin association of the CPC activates Haspin and Bub1, which in turn phosphorylate H3 Thr3 and H2A Thr120, respectively. These histone phosphorylation marks increase the affinity of CPC for chromatin due to Survivin BIR interaction with H3T3ph [1–3] and possibly binding of Borealin to Sgo1 [1, 21]. This facilitates CPC enrichment at inner centromeres during prometaphase and metaphase. In the future, it will be important to understand how such extensive multivalent interactions between CPC and chromatin are weakened to transfer the CPC from chromatin to the central spindle during late metaphase/anaphase [28, 29].

## Acknowledgements

We thank the staff of the Edinburgh Protein Production Facility and the Centre Optical Instrumentation Laboratory for their help. We also thank Tony Ly, Dhanya Cheerambathur and Andrew Goryachev for critical reading of the manuscript. The Wellcome Trust generously supported this work through a Senior Research Fellowship (202811) to AAJ, a Principal Research Fellowship to WCE (073915), a Senior Research Fellowship to JR (084229), a Sir Henry Dale Fellowship to PV (104175/Z/14/Z), a Centre Core Grant (092076 and 203149) and an instrument grant (091020) to the Wellcome Trust Centre for Cell Biology, a Multi-User Equipment grant 101527/Z/13/Z to the EPPF. JGR was supported by the Marie Curie Action PloidyNet, funded by the European Union Seventh Framework Programme (FP7/2007–2013) under Grant Agreement Number 607722.

## STAR Methods

### Protein Expression and purification of the CPC

Survivin was cloned as a 3C-cleavable His-GFP tagged protein in a pRSET vector (Thermo Fisher Scientific). The different Borealin fragments were cloned as a TEV-cleavable His-tagged protein in a pETM vector (gift from C. Romier, IGBMC, Strasbourg), and INCENP_1-58_ was cloned as an untagged protein in a pMCNcs vector.

The CPC subunits were co-expressed in BL21(DE) pLysS strain. Cells were lysed in lysis buffer (25 mM Hepes pH 7.5, 500 mM NaCl, 25 mM Imidazole, 2 mM β-mercaptoethanol) and purified using a 5 ml HisTrap HP column (GE Healthcare). The protein-bound column was washed with lysis buffer, followed by wash buffer (25 mM Hepes pH 7.5, 1 M NaCl, 50 mM KCl, 10 mM MgCl, 25 mM Imidazole, 2 mM ATP, 2 mM β-mercaptoethanol). Elution buffer (25 mM Hepes pH 7.5, 500 mM NaCl, 500 mM Imidazole, 2 mM β-mercaptoethanol) was used to elute the proteins. Tags were cleaved overnight with 3C and TEV proteases while dialysing against 25 mM Hepes pH 7.5, 150 mM NaCl, 4 mM dithiothreitol (DTT) at 4°C. The complex was further purified by a cation exchange chromatography (HiTrap SP, GE Healthcare) followed by gel filtration using a Superdex 200 increase 10/300 column (GE Healthcare) pre-equilibrated with 25 mM Hepes pH 8, 200 mM NaCl, 4 mM DTT.

### Expression and purification of recombinant histones and refolding of histone octamers

Human H2A and H2B and *Xenopus laevis* H3 and H4 were purified as described before [32]. Briefly, H2A, H2B and H3 were expressed in BL21 (DE3) pLysS cells while H4 were expressed in BL21 cells using LB media. The histones were purified from inclusion bodies using a Dounce glass/glass homogenizer. After solubilisation of the inclusion bodies we performed a the three-step dialysis against urea dialysis buffer (7 M Urea, 100 mM NaCl, 10 mM Tris pH 8, 1 mM EDTA, 5 mM β-mercaptoethanol). The sample was then applied to a HiTrap Q anion exchange column and a HiTrap SP cation exchange column (GE Healthcare). Histones were eluted from the HiTrap SP column using a linear gradient from 100 mM to 1 M NaCl in 7 M Urea, 10 mM Tris pH 8, 1 mM EDTA and 1 mM DTT. Purified recombinant histones were dialysed against water containing 5 mM β-mercaptoethanol before lyophilization and storage at −80°C.

To generate histone H3 phosphorylated at threonine 3 by native chemical ligation, histone H3 lacking residues 1–31 containing a threonine-to-cysteine substitution at position 32 and a cysteine-to-alanine substitution at position 110 (H3Δ1–31 MT32C C110A) was expressed and purified as described above. Native chemical ligation reactions with H3Δ1–31 MT32C C110A and the N-terminal H3 peptide ARTPhKQTARKSTGGKAPRKQLATKAARKSAPA containing a C-terminal benzyl thioester (Peptide Protein Research Ltd., Fareham, UK) were carried out in 6 M Guanidine HCl, 250 mM sodium phosphate buffer pH 7.2, 150 mM 4-mercaptophenylacetic acid (MPAA), 50 mM TCEP for 72 h at room temperature with constant agitation. Reactions were dialysed three times against 7 M urea, 100 mM NaCl, 10 mM Tris pH 8, 1 mM EDTA, 1 mM DTT. Ligated full-length H3T3Ph histone was separated from unligated truncated histone through cation exchange chromatography on a monoS column (GE) and then dialysed against water containing 5 mM β-mercaptoethanol before lyophilization and storage at −80°C.

Histone octamers were obtained as previously described [32]. Briefly, lyophilized histones were resuspended in unfolding buffer (7M Guanidine HCl, 20 mM Tris pH 7.5, 10 mM DTT) and mixed to equimolar ratios. The histone mix was then dialysed three times against 500ml of refolding buffer (10 mM Tris pH 8, 2M NaCl, 1 mM EDTA, 5 mM β-mercaptoethanol). The octamers were obtained by running the histone mix on a size exclusion chromatography column (Superdex 200 increase 10/300, GE Healthcare) pre-equilibrated with refolding buffer and stored at −80 °C.

### Nucleosome core particle reconstitution

A pBS-601 Widom vector was used to amplify the 147bp 601 Widom positioning sequence with unlabeled, 5’ IR700- or biotin-labelled primers. Mononucleosomes were obtained by using the salt gradient dialysis method [32]. After optimization of the octamer-DNA ratio, the histone octamer-DNA mix was dialysed against TE buffer (10 mM Tris pH 8, 1 mM EDTA, 50 mM NaCl) overnight by gradually decreasing the ionic strength from 2 M using a peristaltic pump.

### Electrophoretic Mobility Shift Assay

Different concentrations of recombinant CPC were added to 20 nM IR700-labelled NCPs in reaction buffer (10 mM Tris pH 7.5, 100 mM NaCl, 1mM MgCl_2_, 1 mM DTT, 1% Glycerol, 0.1mg/ml BSA). Reactions were incubated 1 h at 4 °C and run in a 6 % polyacrylamide native gel in 0.5X TBE buffer at 100V for 2h at 4°C. The fluorescent bound and unbound NCPs were detected with Odyssey CLx Infrared Imaging System (LI-COR Biosciences). The fluorescent signal of the band corresponding to unbound NCPs was quantified using Image J. Values were plotted and statistically analysed using Prism 6.0 (GraphPad Software, Inc).

### Surface Plasmon Resonance

Surface plasmon resonance experiments were performed using a BIAcore T200 instrument (GE Healthcare). Strepavidin coated sensor surfaces (Sensor Chip SA; GE Healthcare) were primed prior to ligand immobilization by three sequential 30 second-injections of 50mM NaOH; 1M NaCl followed by 180 seconds of Running Buffer (25mM Hepes, 250 mM NaCl, 1 mM DTT and 0.05% Tween-20, pH8) at 30 µl/min. Biotinylated ligands (*in vitro* reconstituted NCPs and DNA) were then immobilized on respective flow-cells by injecting 20 nM solutions of respective ligands at 5 µl/min, and varying the contact time until the Response Units (RU) on the surface reached ∼ 500 RU. Immediately prior to each SPR experiment, samples (CPC complexes, NCPs and DNA) were dialysed against running buffer for 1h at 4 °C. Two-fold dilution series, of respective analytes, from 7.8 nM – 1 µM in Running Buffer, were injected over the sensor surface at 100 µl/min at 8 °C, with 90 second association and 150 second dissociation times. This was then immediately followed by a 300 second regeneration phase in Running Buffer. A neutravidin surface without ligand served as a reference flow-cell for bulk correction. Apparent equilibrium dissociation constants were calculated from the sensorgrams by global fitting of a steady state, 1:1 binding model, with mass transport considerations, using the analysis software (v2.02), provided with the BIAcore T200 instrument. Data was replotted for clarity using Prism 6.0 (GraphPad Software Inc).

### Rescue experiments and Immunofluorescence Microscopy

Depletion of endogenous Borealin using RNAi and rescue experiments were performed as previously described [9] using jetPRIME (Polyplus Transfection). HeLa cells were grown on coverslips in twelve-well plates and medium was changed 12 h after transfection. Cells were fixed in 4 % paraformaldehyde (PFA) 36 h after transfection. The oligonucleotides targeting the 3’ UTR of Borealin (5’-AGGTAGAGCTGTCTGTTCAdTdT-3’) [9] or targeting luciferase as a control (5’-CGUACGCGGAAUACUUCGAdTdT-3’) [33], were described previously.

For quantification of the Survivin signal, HeLa CDK1 analogue sensitive (CDK1-as) cells were used [27]. Cells were synchronised for 14 h using 10 µM 1NM-PP1 and fixed 90 min after washout to increase the number of cells in metaphase.

RNAi depletion of Haspin was performed using oligonucleotides described previously [34] (siRNA ID 1093). Cells were transfected using jetPRIME and fixed in 4 % PFA 48 h after transfection.

Following antibodies were used for indirect immunofluorescence: Anti-myc (1:200; 9E10; Merck Millipore), anti-Borealin (1:500; 147-3; MBL), anti-Survivin (1:500; NB500-201; Novus), anti-H3T3ph (1:500; 07-424; Upstate), anti-H2AT120ph (1:500, 61195, Active Motif), anti-ACA (1:300; 15-235; Antibodies Inc). Hoechst 33342 was used for DNA staining. Imaging was performed using a wide-field DeltaVision Spectris (Applied Precision) microscope with a 100x NA 1.4 PlanApo objective. Shown images are maximum intensity projections. The acquired images were deconvolved using SoftWoRx 3.6 (Applied Precision) and the centromere intensity of Survivin was quantified using an ImageJ plugin (DOI: 10.5281/zenodo.2574963). Briefly, the plugin quantifies the mean fluorescence signal of Survivin in a 2 pixel-wide ring immediately outside the centromere, defined with the ACA staining. For background subtraction, a selected area within the cytoplasm signal was selected. Statistical significance of the difference between normalised intensities at the centromere region was established by a Mann-Whitney test using Prism 6.0 (GraphPad Software).

### Chemical Crosslinking and MS analysis

Crosslinking experiments of the CPC-NCP complexes were performed using 1-ethyl-3-(3-dimethylaminopropyl) carbodiimide (EDC, Thermo Fisher Scientific) in the presence of N-hydroxysulphosuccinimide (NHS, Thermo Fisher Scientific). EDC is a zero-length chemical crosslinker capable of covalently linking primary amines of Lysine and the protein N-terminus and to a lesser extend also hydroxyl groups of Serine, Threonine and Tyrosine with carboxyl groups of Aspartate/Glutamate. 8 µg of CPC or CPC-NCP complexes were incubated with 30 µg EDC and 66 µg of N-hydroxysulphosuccinimide for 2 hours at room temperature. The crosslinking was stopped by the addition of 100 mM Tris-Cl. Crosslinking products were resolved using 4-12 % Bis-Tris NuPAGE (Invitrogen) for 5 min and briefly stained using Instant Blue (Expedeon). Bands were excised, and the proteins were reduced with 10 mM DTT for 30 min at room temperature, alkylated with 55 mM iodoacetamide for 20 min at room temperature and digested using 13 ng/µl trypsin (Promega) overnight at 37°C. The digested peptides were loaded onto C18-Stage-tips [35] for LC-MS/MS analysis. LC-MS/MS analysis was performed using Orbitrap Fusion Lumos (Thermo Fisher Scientific) with a “high/high” acquisition strategy. The peptide separation was carried out on an EASY-Spray column (50 cm × 75 μm i.d., PepMap C18, 2 μm particles, 100 Å pore size, Thermo Fisher Scientific). Mobile phase A consisted of water and 0.1% v/v formic acid. Mobile phase B consisted of 80% v/v acetonitrile and 0.1% v/v formic acid. Peptides were loaded at a flow rate of 0.3 μl/min and eluted at 0.2 μl/min using a linear gradient going from 2% mobile phase B to 40% mobile phase B over 109 or 139 min (each sample has been running three time with different gradients), followed by a linear increase from 40% to 95% mobile phase B in 11 min. The eluted peptides were directly introduced into the mass spectrometer. MS data were acquired in the data-dependent mode with 3 s acquisition cycle. Precursor spectra were recorded in the Orbitrap with a resolution of 120,000. The ions with a precursor charge state between 3+ and 8+ were isolated with a window size of 1.6 m/z and fragmented using high-energy collision dissociation (HCD) with collision energy 30. The fragmentation spectra were recorded in the Orbitrap with a resolution of 15,000. Dynamic exclusion was enabled with single repeat count and 60 s exclusion duration. The mass spectrometric raw files were processed into peak lists using ProteoWizard (version 3.0.6618) [36], and cross-linked peptides were matched to spectra using Xi software (version 1.6.743) [37] with in-search assignment of monoisotopic peaks [38]. Search parameters were MS accuracy, 3 ppm; MS/MS accuracy, 10ppm; enzyme, trypsin; cross-linker, EDC; max missed cleavages, 4; missing mono-isotopic peaks, 2; fixed modification, carbamidomethylation on cysteine; variable modifications, oxidation on methionine and phosphorylation on threonine for phosphorylated sample; fragments, b and y ions with loss of H2O, NH3 and CH3SOH. FDR was computed using XiFDR and results reported at 5% residue level FDR [39].

