## Supplemental Data for "Direct Nucleosome Binding of Borealin Secures Chromosome Association and Function of the Chromosomal Passenger complex"

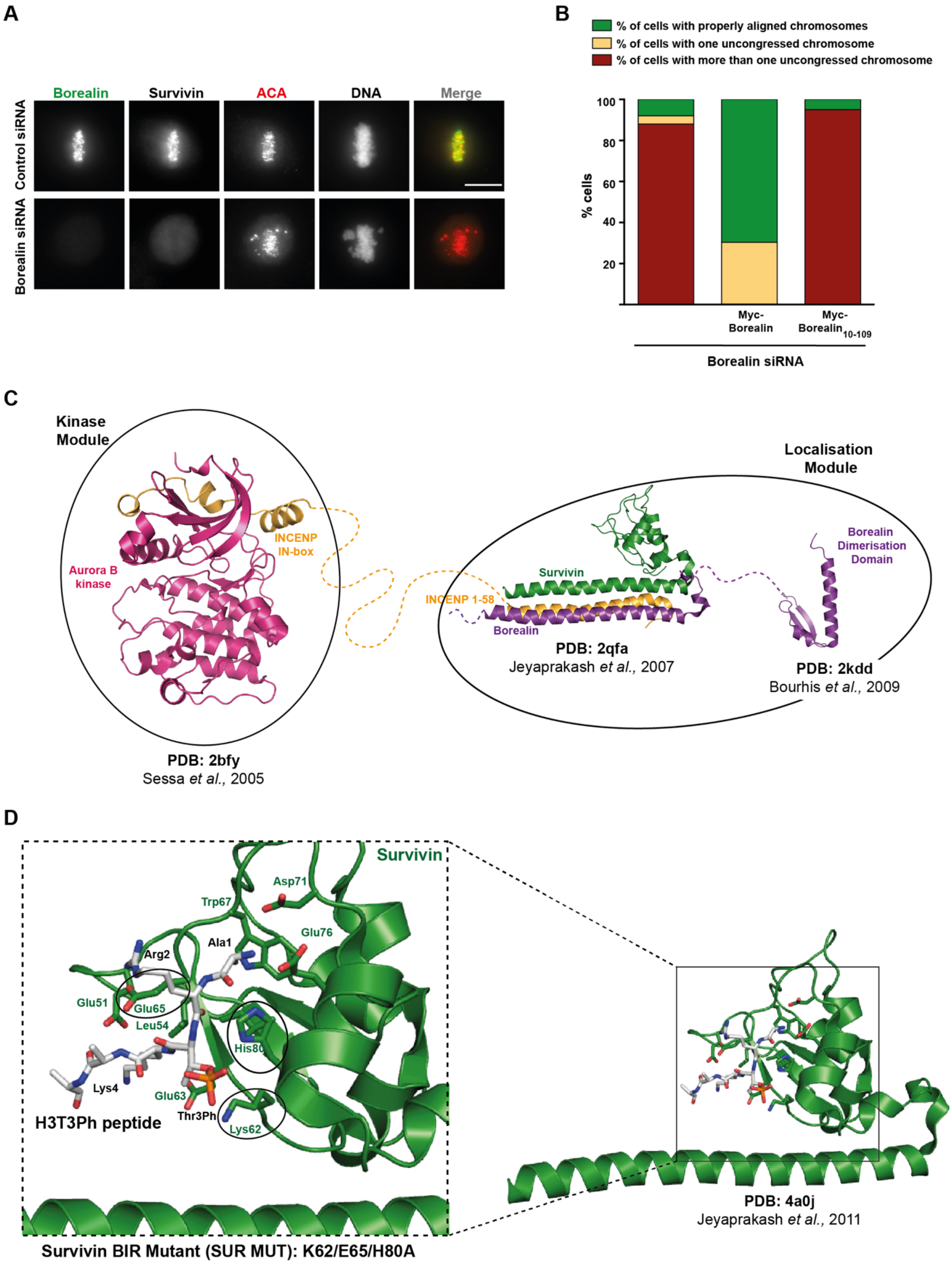


**Figure S1. Structural organisation of the Chromosomal Passenger Complex.**

(A) Treatment of siRNA oligos targeting Borealin 3'UTR resulted in no Borealin signal from early prophase to anaphase. Immunofluorescence staining of Borealin, Survivin and ACA in

HeLa cells. Hoechst was used for DNA staining. Images were taken 36 h after transfection. Scale bar, 10 μm. (B) Analysis of cells showing uncongressed chromosomes of the siRNA-rescue assay for Myc-Borealin$_{10-109}$ fragment shown in Fig. 1B. A total of 25, 23 and 20 cells were counted for Borealin siRNA, Myc-Borealin and Myc-Borealin$_{10-109}$ rescue conditions, respectively. (C) The CPC can be divided into a localisation module (composed of Borealin, Survivin and INCENP 1-58) and a kinase module (composed of Aurora B and INCENP IN-box) connected by a central helical coiled coil of INCENP. The structures have been generated using Pymol v2.0.6. (D) Close-up view of the histone H3 peptide with a phosphorylated Thr3 (grey) bound to Survivin (green). Survivin residues involved in H3 tail binding are shown in stick representation. Amino acid residues mutated in the BIR domain of Survivin (SUR MUT: K62/E65/H80A) to abolish H3-tail binding are highlighted in circles.

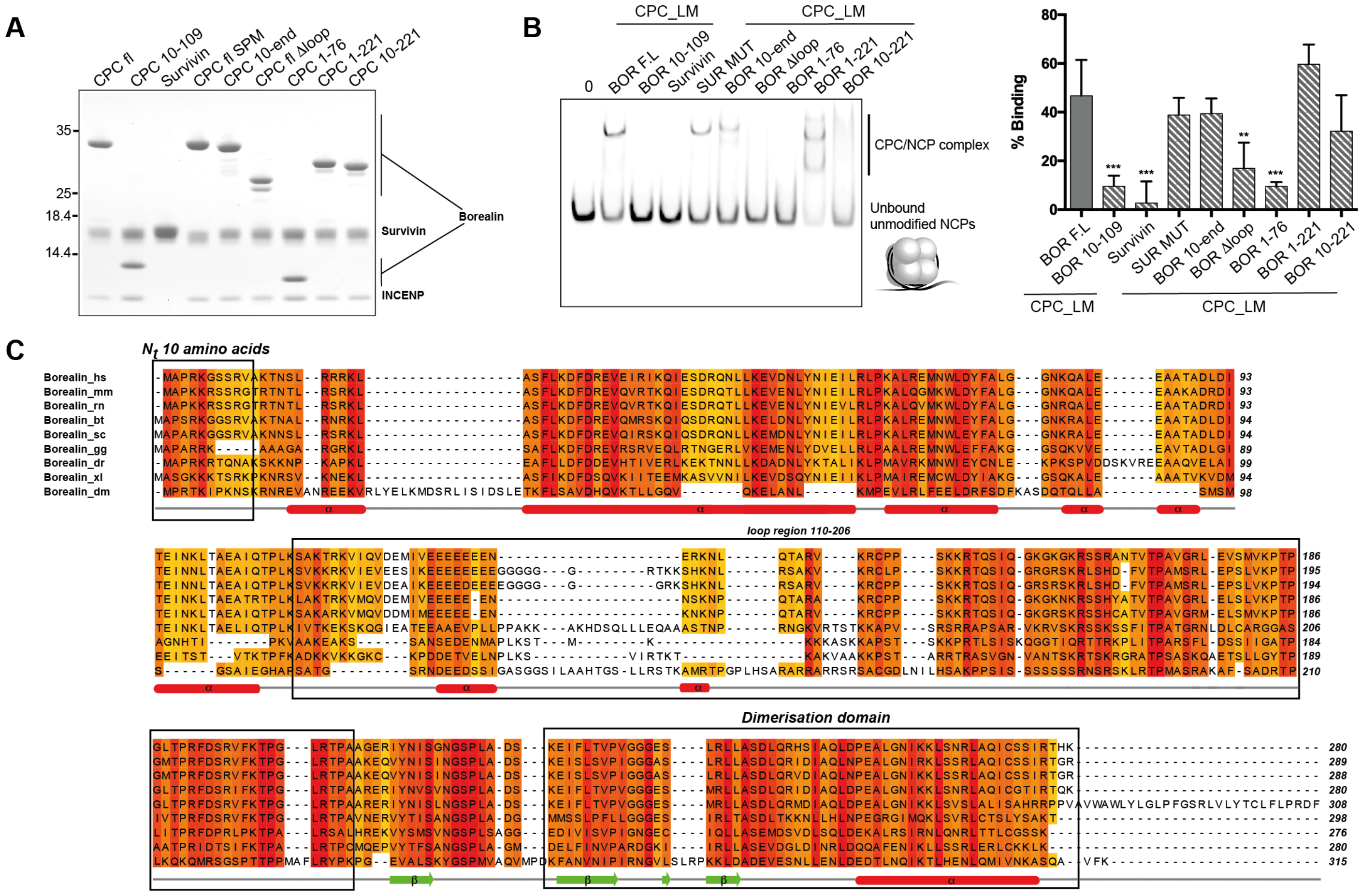


**Figure S2. Multiple regions of Borealin contribute to nucleosome binding**

(A) Representative Coomassie-stained SDS-PAGE analysis of purified recombinant CPC complexes used in the EMSA and SPR assays. (B) Native PAGE analysis of an EMSA assay performed with recombinant CPC_LM complexes containing different Borealin truncations to evaluate their binding to unmodified IR700-labelled NCPs (left) and quantification of binding (right). Concentrations of NCP and CPC used in the assay were 20 nM and 160 nM, respectively. Mean of % of binding ± standard deviation (SD); n=5; **P ≤ 0.01, *** P ≤ 0.001, unpaired *t*-test. (C) Amino acid conservation of Borealin (conservation score is mapped from red, highly conserved, to yellow, poorly conserved). The alignment includes Borealin orthologs from *H. sapiens* (hs), *Mus Musculus* (mm), *Rattus norvegicus* (rn), *Bos Taurus* (bt), *Sus scrofa* (ss), *Gallus gallus* (gg), *Danio rerio* (dr), *Xenopus laevis* (xl), *Drosophila* *melanogaster* (dm). Secondary structure elements are depicted below the sequence alignment. The black box highlights regions we show here critical for nucleosome binding. Multiple sequence alignment was performed with Clustal Omega (EMBL-EBI) and edited with Jalview 2.10.5 [1].

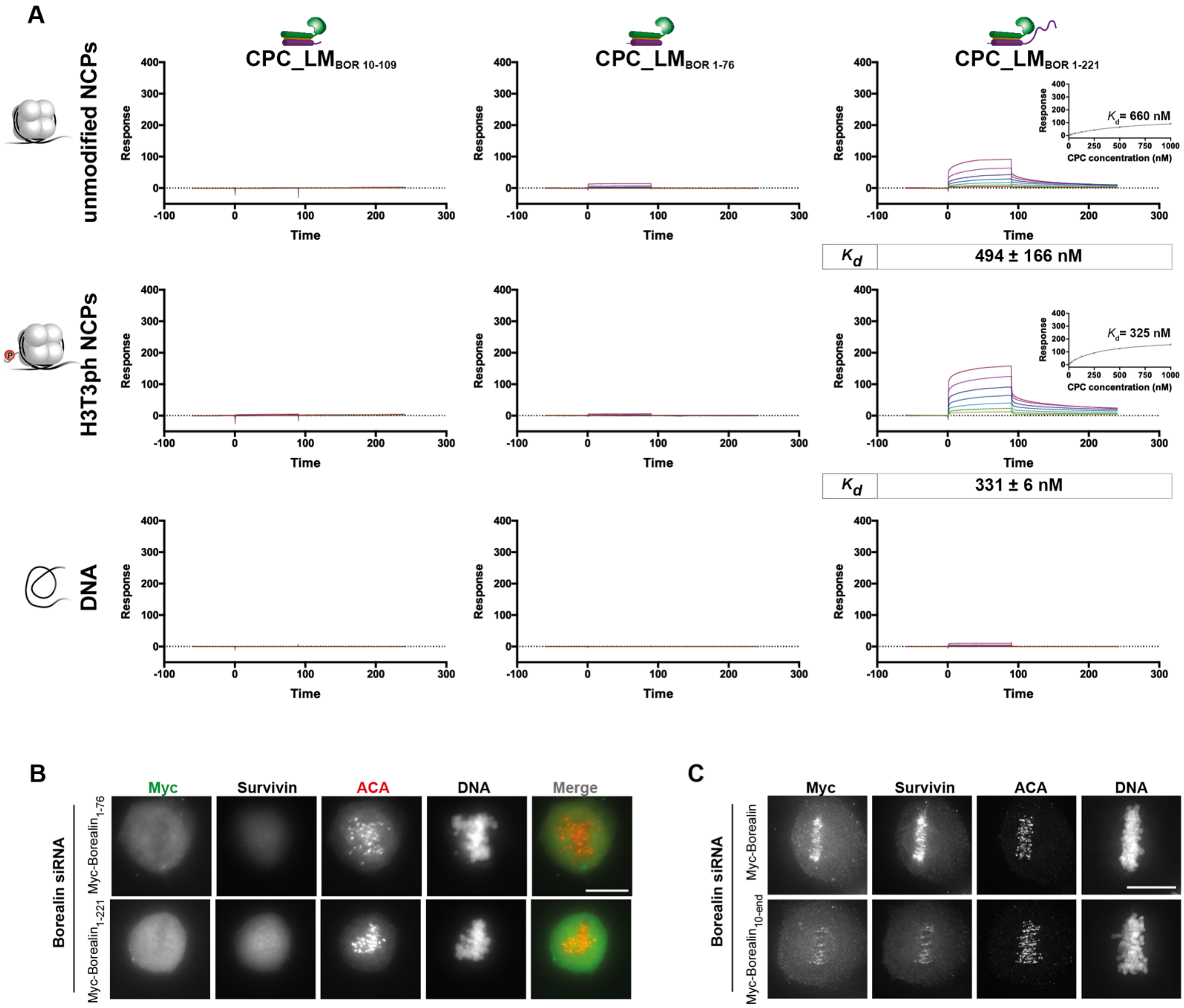


**Figure S3. Biophysical and cellular characterisation of Borealin mutants in facilitating nucleosome binding and chromosome association of CPC.**

(A) Representative SPR sensorgrams of the interaction between CPC_LM complexes containing different Borealin C-terminal truncations (CPC_LM$_{BOR\ 10\text{-}109}$, CPC_LM$_{BOR\ 1\text{-}76}$ and CPC_LM$_{BOR\ 1\text{-}221}$) and unmodified or H3T3ph NPCs or DNA immobilized on the surface of a neutravidin sensor chip. Mean values (n ≥ 2, ± SEM) determined for the equilibrium dissociation constant ($K_d$) are shown. (B) Representative fluorescence images for the evaluation of a siRNA-rescue assays for Borealin$_{1\text{-}76}$, Borealin$_{1\text{-}221}$ constructs.

Immunofluorescent staining of Myc, Survivin and ACA in HeLa cells co-transfected with siRNA duplexes targeting the 3'UTR region of Borealin and Myc-Borealin constructs. Hoechst used for DNA staining. Scale bar, 10 μm. (C) Representative fluorescence images used for the quantification of centromere levels of Survivin shown in Fig. 2D. Images were deconvolved using SoftWoRx 3.6.

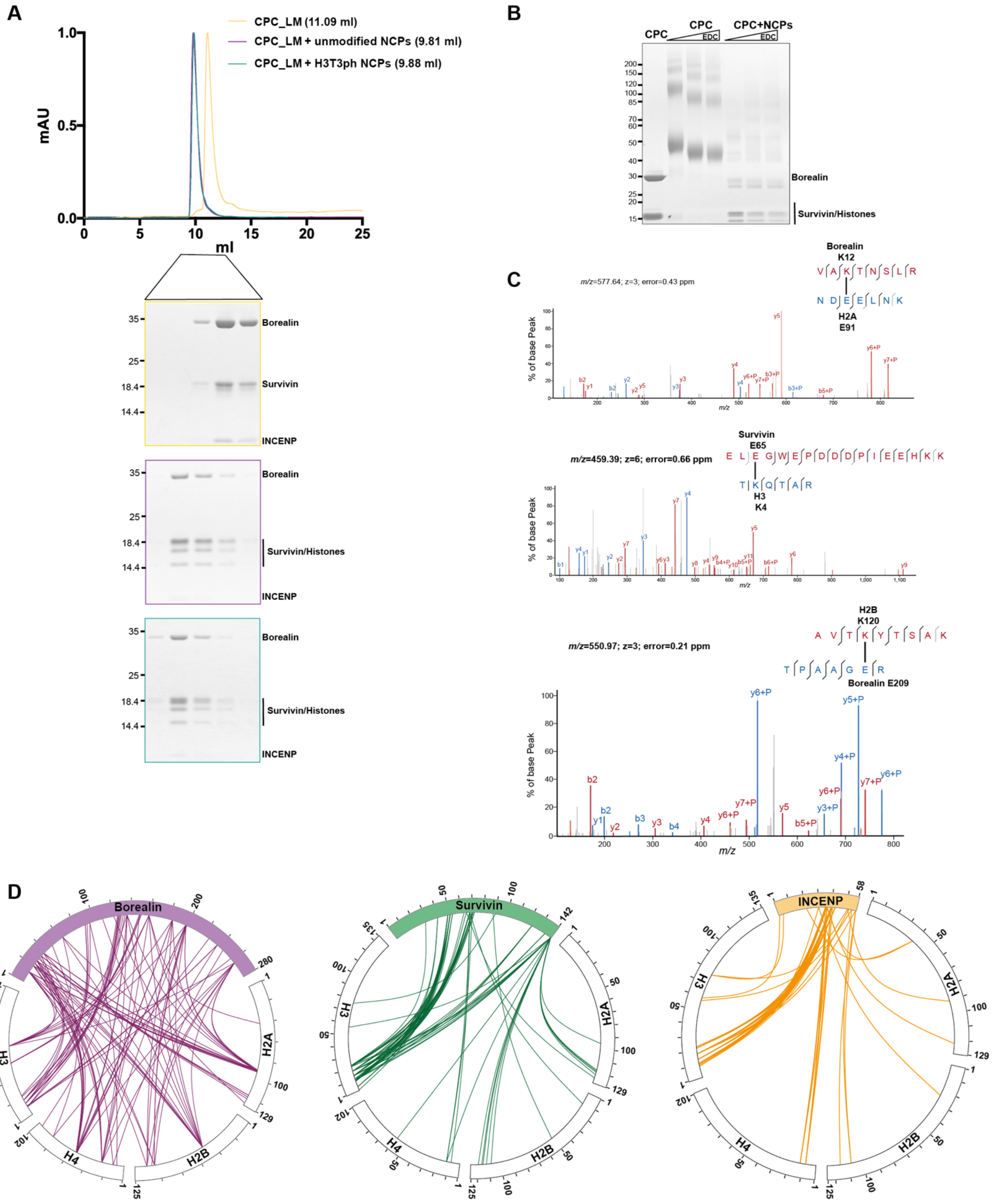


**Figure S4. Mapping of CPC-NCP interactions using chemical crosslinking and mass spectrometric analysis.**

(A) Size exclusion chromatogram of recombinant CPC_LM (yellow) and CPC_LM in complex with unmodified (magenta) or H3T3ph NCPs (cyan) using a Superdex 200 increase 10/300 GL column (top). Coomassie-stained SDS-PAGE analysis of samples resolved in the Superdex 200 column (bottom). (B) Representative SDS-PAGE of 8 μg of CPC or CPC-NCP complex crosslinked with EDC crosslinker. (C) High resolution fragmentation spectra of a few representative crosslinked peptides, displayed using XiSpec (Ref: PMID: 29741719). (D) Crosslink mapping of interactions between the CPC subunits (Borealin-purple, Survivin-green, INCENP-yellow) and histones from H3T3ph NCPs. Intermolecular contacts involving Borealin, Survivin and INCENP and histones are shown as purple, green or yellow lines.

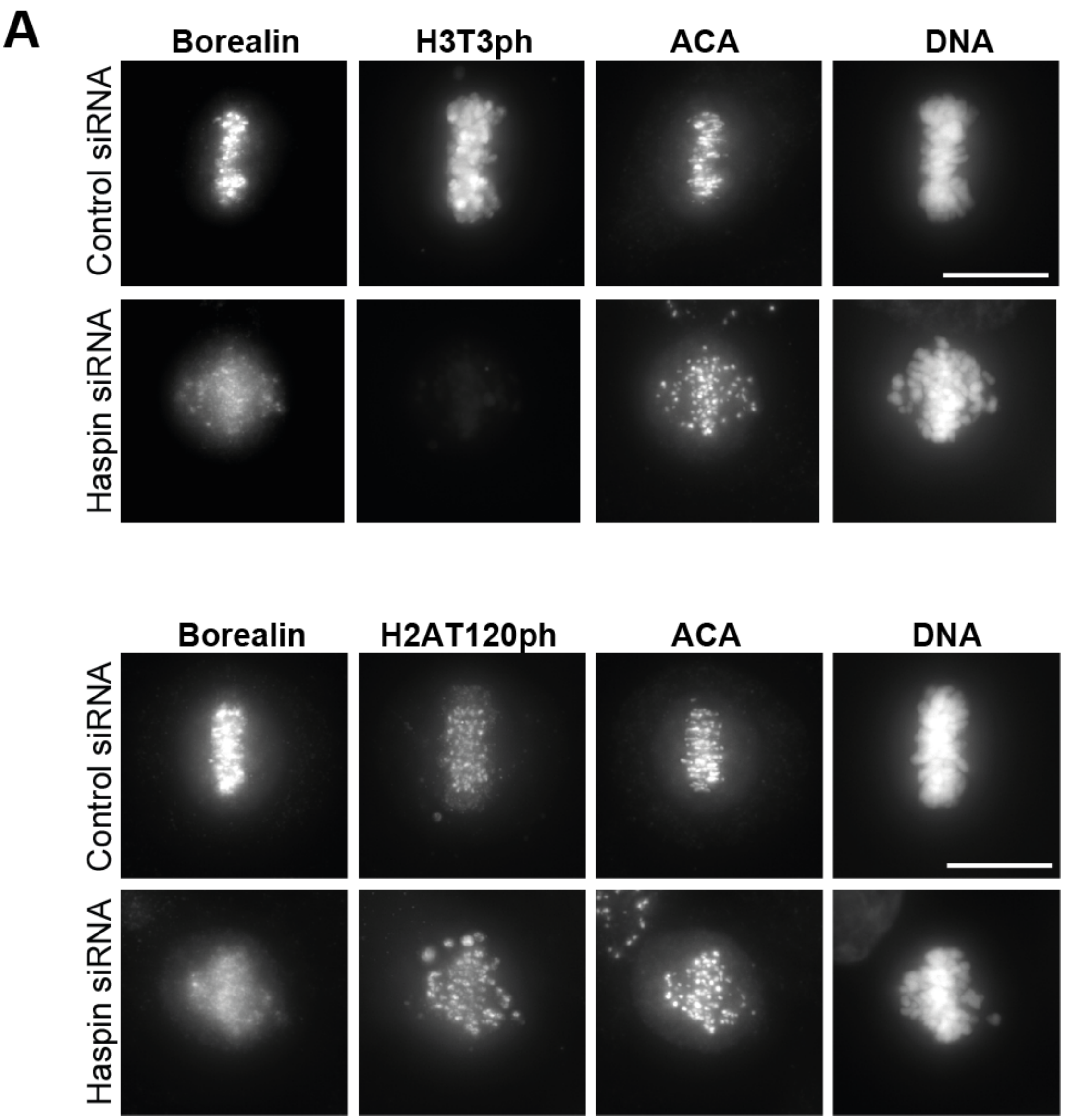


**Figure S5. Bub1-mediated H2A T120 phosphorylation does not depend on Haspin activity.**

(A) Immunofluorescence analysis of Borealin, H3T3ph, H2AT120ph and ACA in HeLa cells transfected with siRNA duplexes targeting the Haspin transcript for 48 h and Control siRNA duplexes. Hoechst staining was used to visualise DNA.
